# Identification of multivariate phenotypes most influenced by mutation: *Drosophila serrata* wings as a case study

**DOI:** 10.1101/2025.02.25.640174

**Authors:** Cara Conradsen, Katrina McGuigan

**Affiliations:** School of the Environment, The University of Queensland. St. Lucia, QLD, 4072 Australia; European Centre for Environment and Human Health, University of Exeter Medical School, Environment and Sustainability Institute, Treliever Road, Penryn Campus, Penryn, TR10 9FE, United Kingdom; Environment and Sustainability Institute, University of Exeter, Treliever Road, Penryn Campus, Penryn, TR10 9FE, United Kingdom

**Keywords:** pleiotropy mutational variance, mutational covariance, DsGRP ential

## Abstract

The distribution of pleiotropic mutational effects impacts phenotypic adaptation. However, small effect sizes and high sampling error of covariances hinder investigations of the factors influencing this distribution. Here, we explored the potential for shared information across traits affected by the same mutations to counter sampling error, allowing robust characterisation of patterns of mutational input. Exploiting a published dataset representing 12 samples of the same mutation accumulation experiment in *Drosophila serrata*, we inferred robust signals of mutational effects from the concordance across samples. Krzanowski common subspace analysis identified a multivariate wing trait with statistically supported mutational variance in all samples. Importantly, this multivariate trait was aligned with the major axis of among-line (mutational) variance within most population samples. That is, despite considerable heterogeneity among samples in individual (co)variance parameter estimates, the predominant pattern of correlated mutational effects was identified in datasets reflecting a typical mutation accumulation experimental design. Two other multivariate traits were statistically supported across most samples. Smaller effect sizes (lower mutational variance) with concomitant larger sampling error or other factors (e.g., microenvironmental dependence of effects) may reduce the robustness of estimated mutational input for these traits. Overall, our results suggest sampling error does not preclude multivariate analyses of mutation accumulation experiments from extending our knowledge of pleiotropic mutational effects.

## INTRODUCTION

How spontaneous mutations affect the phenotype is central to a range of evolutionary genetic phenomena, including the evolutionary dynamics of genetic variance and the evolutionary fates of populations (Johnson & Barton, 2005; Schultz & Lynch, 1997; Whitlock, 2000; Zhang & Hill, 2002). Reflecting this broad significance, substantial effort has been devoted to characterising the input of new mutation to individual quantitative traits (Conradsen et al., 2022; Halligan & Keightley, 2009; Houle et al., 1996; Walsh & Lynch, 2018). However, this individual trait focus is at odds with the long-standing recognition that selection does not act on individual traits in isolation (Svensson et al., 2021) and that the shared genetic basis of traits influences their response to selection (Walsh & Blows, 2009). Furthermore, theory suggests the strength of correlation of mutational effects across traits is a key determinant of the stability of quantitative genetic covariances, and thus of our ability to predict evolution over multi-generational timescales (Jones et al., 2003).

Several studies have extended beyond individual traits to consider the bivariate distribution of mutational effects, as reflected in the magnitude of trait correlation arising due to novel mutation. Studies estimating mutational correlations between broadly defined reproductive output and survival traits suggest mutational effects on these traits may typically be positively correlated (median mutational correlation = 0.33: Fernandez & Lopez-Fanjul, 1996; Houle et al., 1994; Keightley et al., 2000; Keightley & Ohnishi, 1998; Martorell et al., 1998). Nonetheless, correlations between such traits vary widely in magnitude (−0.5 to 1), and estimates have large confidence intervals. Mutational correlations between morphological, behavioural, locomotor or metabolic traits have also been reported, with estimates spanning moderately negative through strongly positive (e.g., Clark et al., 1995; Estes et al., 2005; Keightley & Ohnishi, 1998; Mackay et al., 1992; Santiago et al., 1992). Where the same traits have been considered in different studies, as with reproduction and survival traits, marked differences in the mutational correlation estimates are observed (e.g., activity of specific metabolic enzymes: Clark et al., 1995; Harada, 1995). It remains unresolved whether heterogeneity among mutational correlation estimates reflects evolutionarily meaningful differences — such as due to genetic background, taxon or trait type — or instead arises from methodological inconsistencies, differences in trait definitions, or the inherent estimation error associated with quantitative genetic parameters.

As pairwise correlations may offer limited insight into the evolution of phenotypes or genetic variance (Blows & McGuigan, 2015; Johnson & Barton, 2005; Svensson et al., 2021; Walsh & Blows, 2009), it is also important to adopt a broader multivariate (geometric) approach in studies of mutational covariance across larger trait sets. Several more recent studies have interrogated the higher dimensional patterns of correlation in mutational effects captured by the mutational variance covariance matrices (**M**). Intriguingly, some of these studies have found evidence of concordance in major axes between **M** and a corresponding standing genetic variance covariance matrix (**G**) (Dugand et al., 2021; Houle et al., 2017), while others have found marked differentiation between **M** and **G** (Latimer et al., 2014; François Mallard et al., 2023). While further empirical data is essential to clarify the evolutionary implications of these contrasting observations, it is equally critical to improve our understanding of the statistical properties of **M**—especially the sources and consequences of estimation error.

High estimation error is a well-known challenge to studying quantitative genetic parameters (Klein, 1974; Klein et al., 1973; Lynch & Walsh, 1998). Particularly when the number of independent samples (e.g., mutation accumulation lines) is small relative to the number of variables (traits), random sampling error can generate extensive spurious covariance, leading to inflation of the estimated magnitude of variance along leading eigenvectors of covariance matrices (Blows & McGuigan, 2015; Johnstone, 2007; Sztepanacz & Blows, 2017). The potential influence of sampling error on estimates and conclusions is likely to be exacerbated for mutational relative to standing genetic variation due to the smaller magnitude of (co)variances (i.e., a smaller statistical effect size). For individual traits, mutations contribute, on average, 0.24% to the phenotypic variance per generation (Conradsen et al., 2022; Houle et al., 1996; Lynch et al., 1999), in contrast to the median portion of phenotypic variance due to standing genetic variation, which is >10% across almost any trait type (Hansen et al., 2011). Allowing mutations to accumulate over hundreds of generations of mutation-drift evolution can mitigate the difficulty posed by small magnitude (co)variance (Johnson & Barton, 2005; Lynch et al., 1999). However, estimates from such experiments are increasingly likely to reflect the combined effects of multiple mutations per genotype (mutation accumulation line), which will limit insight into the distribution of effects of individual mutations. Furthermore, logistical limitations might lead to long-term maintenance trading-off with the number of mutationally divergent genotypes (lines) that can be maintained and assayed, opposing the statistical gain from sampling more mutations per genotype.

These concerns about statistical limits may disincentivise investment in the further empirical estimates of **M** that are required to advance our understanding of this important evolutionary parameter. However, by leveraging shared signal among covarying traits (Schmitz et al. 1998), multivariate analyses may, in fact, offer a powerful solution for advancing our understanding of mutational effect distributions. This strategy has proven effective in human genome-wide association studies, where multi-trait frameworks have significantly improved variant detection across diverse traits (Baselmans et al., 2019; Qi & Chatterjee, 2018; Turley et al., 2018; Wu, 2020). Here, we seek to determine whether shared mutational signal might similarly lead to relatively robust identification of the major axis of mutational variation (i.e., the first eigenvector of **M**, **m**ₘₐₓ).

To address this question, we take advantage of a unique published dataset in which it is possible to repeatedly estimate **M** for the same mutation accumulation experiment. This data represents sequential samples of *Drosophila serrata* wing phenotypes in 42 lines that have diverged through mutation-drift over 20 generations (Conradsen et al., 2021). There is clear evidence from developmental studies (reviewed in Blair, 2007; Matamoro-Vidal et al., 2015), identification of causal loci (e.g., Pitchers et al., 2019) and estimates of both **G** (McGuigan & Blows, 2007; Mezey & Houle, 2005) and **M** (Dugand et al., 2021; Houle et al., 2017) of pleiotropic effects of alleles across Drosophila wing traits. Thus, we expect traits to share mutational information, essentially resulting in higher replication of phenotypic measures and increased precision of estimation. However, it is unclear whether other factors (e.g., developmental error, measurement error, micro-environmental effects) also generate shared, but uninteresting, information, negating the multivariate gain in power.

## METHODS

### Population history and wing data collection

The experimental design was described in detail in Conradsen et al. (2022), where mutational variance of individual traits was investigated, but no consideration was given to mutational covariance among traits. Briefly, a highly inbred *D. serrata* Reference Genome Panel (DsRGP) line (Reddiex et al., 2018), DsGRP-226, was used as a progenitor to establish 200 Mutation Accumulation (MA) lines (Kannan et al., 2023). After 20 generations of mutation accumulation via mutation-drift in brother-sister inbreed lines, 42 MA lines were randomly chosen and each used to establish two new sublines, one under each of two population size treatments (Small and Large: Figure 1a). For each of the 84 sublines (42 MA x 2 treatments), wings were sampled from up to six male flies in each of two replicate rearing vials for six sequential generations (Figure 1a): a total of 5,135 wings were available for analysis, corresponding to 12 (2 treatments by 6 generations) repeated samples in which among-line (mutational) variance could be estimated.

**Figure 1.**
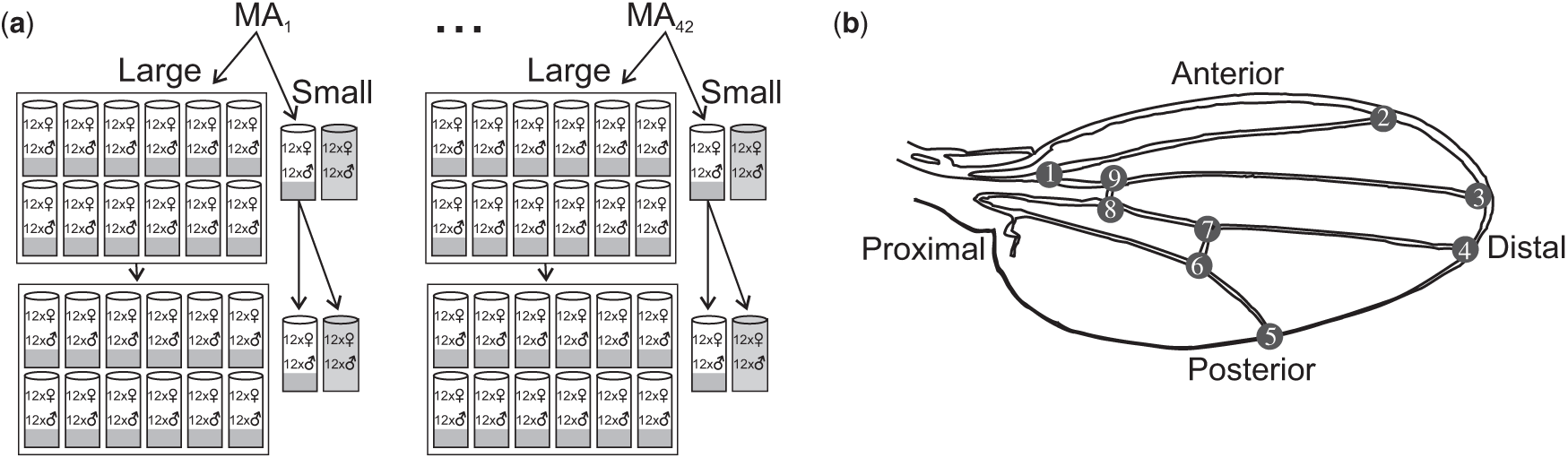
Schematic of (a) experimental design and (b) *Drosophila* wing traits. (a) 42 MA lines each founded two sublines: Small (S; 12 virgin males and 12 virgin females) and Large (L; 144 virgin males and 144 virgin females, distributed evenly among 12 vials). These 84 lines (S and L subline per 42 MA lines) were maintained at these census population sizes for six generations (first two shown here). Each generation, all emergent flies from the 12 vials per L subline were pooled prior to virgin collection. For S sublines, the focal vial contributed offspring to the next generation, while a second, replicate, vial (grey shaded) was used for wing collection only. Each generation, six randomly chosen males were sampled from each of two vials per line (focal and replicate vials for S; randomly chosen two for L). (b) Positions of nine landmarks were recorded and aligned. Six traits were analysed, five inter-landmark distances (named by their end landmarks: ILD1.2, ILD1.5, ILD2.5, ILD2.8, ILD3.7) and wing size, characterised as centroid size (CS).

The population size treatment, and the sequential sampling introduce the opportunity for segregating new mutations and microenvironment to differentially influence among-line variance estimates, including via environment-specific mutational effects. However, Conradsen et al. (2022) partitioned these potential sources of variance, and provided evidence that sampling error alone accounted for the vast majority of the substantial heterogeneity among the 12 estimates of among-line (mutational) variance, with only a small amount of the variance in one trait (wing size) accounted for by micro-environment and mutation-drift-selection processes. Here, we therefore consider these data as repeated measurement of the effects of mutations fixed in lines during the 20 generations of mutation accumulation, and focus our interpretation on the contribution of sampling error to variation among the 12 estimates of mutational covariance parameters.

Nine wing landmark positions (Figure 1b) were recorded from images, aligned using a generalised Procrustes least squares superimposition, and centroid size (CS, the square root of the sum of the squared distances between each land-mark and their centroid) determined (Rohlf, 2007). Inter-landmark distances (ILDs, in units of centroid size) were calculated from the aligned coordinates. Using the model described below, we attempted to estimate the among-line variance-covariance matrix for the 11 traits previously analysed individually by Conradsen et al. (2022). However, this model failed to converge – a likely consequence of the relatively few lines (42) and the small magnitude of among-line variance accumulated after 20 generations of mutation-drift evolution. We therefore report on a subset of those 11 traits, and consider further in the Discussion how our pragmatic, but non-random, trait selection may influence conclusions. Four (ILD1.9, ILD3.8, ILD4.8 and ILD5.6) of the original 11 traits were excluded because at least one of the 12 (two treatments measured in six generations) among-line variance estimates was zero (see Figure 5 of Conradsen et al., 2022), causing model convergence failure. A further trait (ILD4.5) also impacted model convergence (models containing this trait returning non-positive definite asymptotic matrices precluding confidence interval estimation) and was therefore also excluded. A final subset of five ILDs, along with CS (Figure 1b), were retained. To facilitate multivariate model convergence, data for each trait was standardised (mean = 0, standard deviation = 1) experiment-wide (across all generations and treatments). Mahalanobis distance was used to identify multivariate outliers (critical value of 𝜒^!^ = 22.46, df = 6, α = 0.01), which were excluded from all analyses (55 outliers across the 5135 flies, < 1.1%).

### Estimating mutational variance-covariance matrices

We estimated the 12 among-line (**M**) covariance matrices for our six traits using restricted-maximum likelihood (REML). The MIXED procedure in SAS v9.4 (SAS Institute Inc., Cary, NC.) was used to fit:

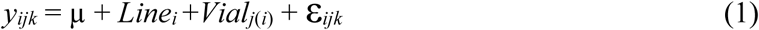

where *y_ijk_* was the vector of six trait values for the *k*th wing (*k* = 1…6), for the *j*th vial (*j* = 1,2), within the *i*^th^ line (*i* = 1…42); µ was the vector of global means per trait; the random effects of *Line*, *Vial* and the residual error (ε) estimated the 6 x 6 unstructured covariance matrices corresponding to among-line (mutational), among replicate vial and among individual flies within a vial, respectively.

To determine the number of statistically supported independent axes of among-line variation, a factor-analytic approach (Hine & Blows, 2006) was applied, using likelihood ratio tests (LRT) to compare sequential models constraining among-line variance from zero up to six dimensions. The LRT degrees of freedom were defined by the difference between models in the number of parameters estimated. Factor analytic modelling conservatively accounts for the effect of sampling error on eigenvalues, representing the most robust approach for determining statistical support for the dimensionality of covariance matrices, (Sztepanacz & Blows, 2017).

To place confidence intervals about estimates we applied the REML-MVN method developed by Meyer and Houle (Houle & Meyer, 2015; 2013), drawing 10,000 random samples from the distribution *N* ∼ (**θ̂, V**), where **θ̂** was the vector of observed (REML) covariance parameter estimates, and **V** was the asymptotic variance-covariance matrix. These RML-MVN samples were used to calculate confidence intervals for both model-estimated and derived parameters (detailed below).

### Constructing null model variance-covariance matrices

The overarching null hypothesis of the current study is that we have 12 repeated measures of the same mutational (co)variance of wing traits, differing from one another only through transient segregation of recent mutations, micro-environmental effects and estimation error. We constructed a null distribution reflecting this by randomly sampling MA lines (with replacement) to create 12 experimental populations (two treatments by six generations) of 42 lines. By retaining the within-line information (i.e., the set of ∼12 flies in replicate vials sampled from a subline each generation were collectively assigned their new, random, population identity) while shuffling lines among population size treatments and generations, we retained the signal of mutation in the populations, focusing our null on the potential for sampling error to generate spurious divergence among 12 repeated measures of **M**. We generated 1000 (randomised) null datasets, and again implemented model 1 to estimate 12 **M** in each of these datasets. Some of the 12,000 null data models failed to converge, resulting in nine of the 1000 null datasets (0.9%) with 11, not 12, estimates of **M**. The null matrices were taken through the same analytical pipeline (detailed below) as the observed data.

### Comparing variance-covariance matrices

We used Krzanowski’s subspace method to determine the extent to which the 12 **M** estimated the same multivariate axes of mutational variance. This approach is described in detail elsewhere (Aguirre et al., 2014; Blows et al., 2004; Krzanowski, 1979). Briefly, R (R Core Team, 2024) was used to diagonalise each **M**, and to estimate the Krzanowski subspace matrix, **H**:

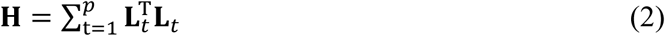

where **L***_t_* contains the subset of the *j*^th^ largest eigenvectors, as rows, of the *t*^th^ **M** matrix, (and 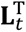 its transpose containing eigenvectors as columns) and *p* = 12 is the number of populations to be compared. Spectral decomposition of **H** describes the degree of shared structure across the matrices, with the eigenvalues ranging from 0 to *p*. Here, an eigenvalue of 12 would indicate that all 12 **M** have variance in the direction of the associated eigenvector of **H**. The interpretation of **H**’s eigenvalues depends on how *j* is defined (Aguirre et al., 2014; Blows et al., 2004; Krzanowski, 1979). When 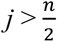 (where *n* is the number of traits; here, *n* = 6, setting *j* > 3), the analysis is constrained to recover common subspaces of two matrices. Conversely, if *j* is allowed to vary among populations, above *j* = min(*j*_1_, *j*_2_, …*j_p_*) dimensions are constrained to be orthogonal (i.e., not contain a common subspace) for at least one population — the one with the lowest *j*. Here, we followed Aguirre et al. (2014) in defining *j* for the *t*^th^ **M** matrix as the number of eigenvectors required to explain at least 90% of the variance; for each of our 12 sampled matrices, this included all dimensions with statistically supported variation (Table 1, Table S2). The 90% criterion ensured that we could interpret at least the first two eigenvalues of **H** (Table 1). For four **M**, *j* = 2, indicating that these matrices were constrained to share no more than two common dimensions with any other matrix. Notably, only one **M** had *j* > 3, indicating that no two matrices were constrained to share common subspaces.

**Table 1.**
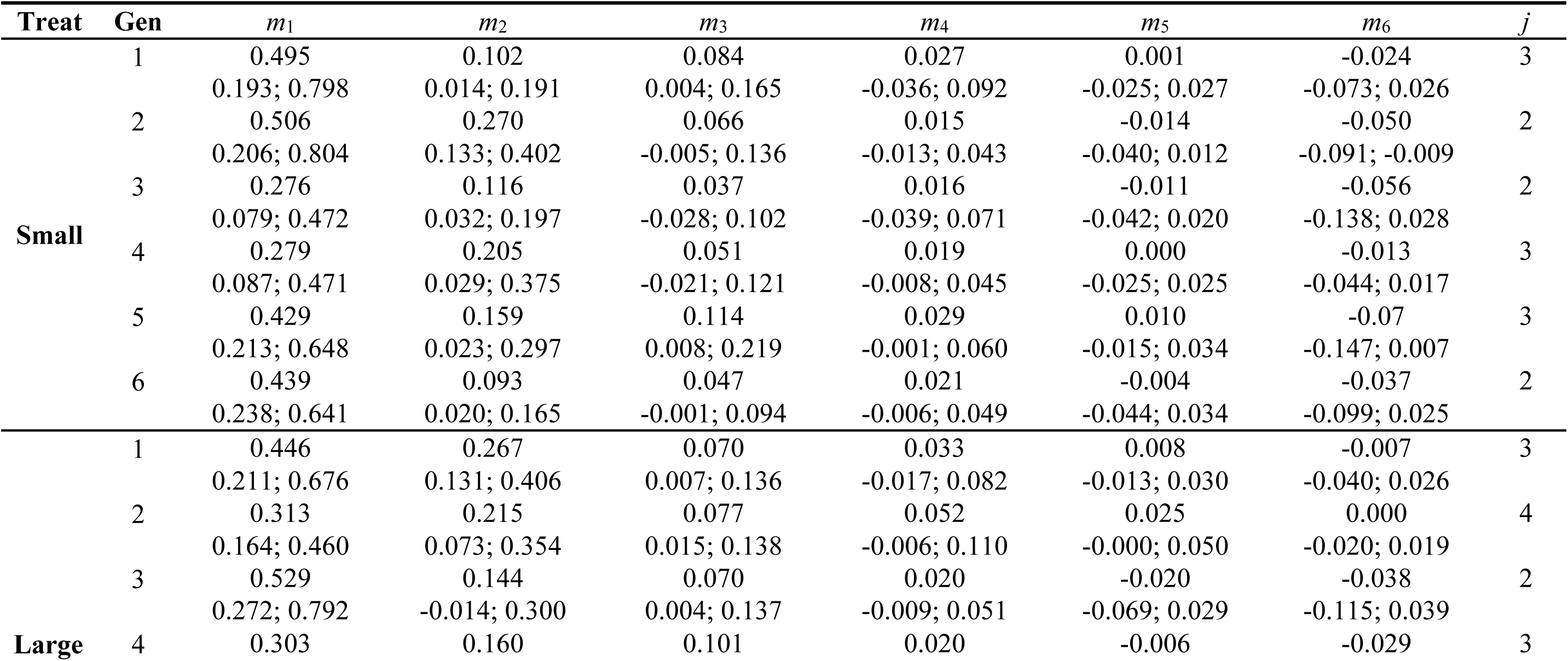
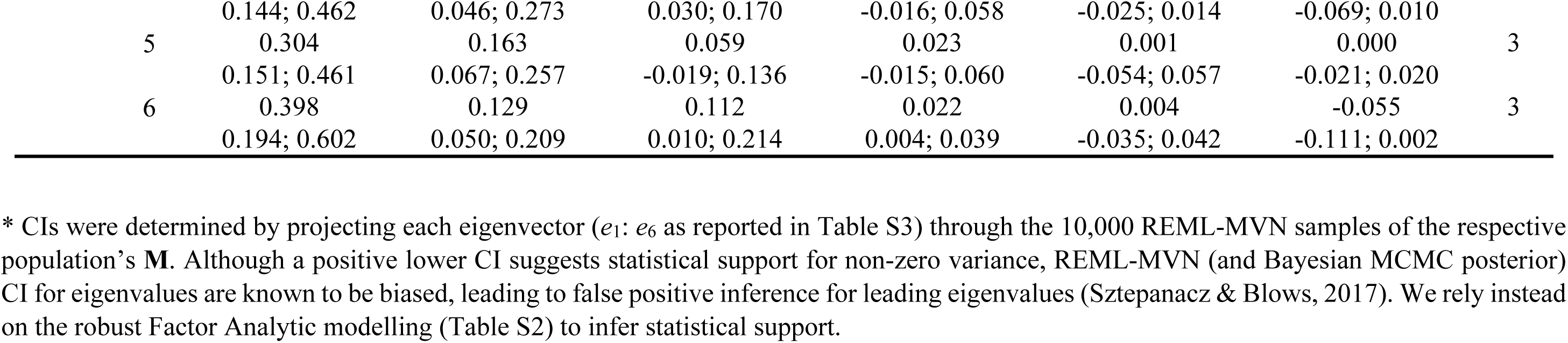
The variance associated with independent axes of trait mutational variance. Eigenanalyses were performed separately for **M** in each population size treatment (Treat: Small or Large) in each generation (Gen: 1 through 6). The eigenvalues (i.e., λ) and their 90% CIs* are reported, along with the number of eigenvectors (*j*) accounting for at least 90% of the variation. These *j* eigenvectors were included in the estimation of the Krzanowski subspace. The estimated **M** and the eigenvector loadings are reported in Tables S1 and S3, respectively.

To place confidence intervals on the eigenvalues of **H**, we applied equation 2, and extracted the eigenvalues of the resulting matrix, to each of the 10,000 REML samples of the 12 matrices. To determine which of the 12 matrices had variance in which parts of the subspace captured by **H**, and whether matrices differed in the magnitude of among-line variance in shared (common) subspaces, we projected the eigenvectors of **H** through each **M** (Aguirre et al., 2014). To place confidence intervals on these estimates of common-subspace variance, eigenvectors of the observed REML **H** matrix were projected through each of the 10,000 REML-MVN samples of the matrix. Equation 2 was also applied to estimate **H** for each of the 1000 null datasets (we used *j* = 3 eigenvectors for all null datasets). The 95% confidence intervals of eigenvalues of the null data **H** were determined directly from distribution across the null 1000 datasets.

## RESULTS

### Among-line variation in the 12 population samples

Consistent with previous analyses of each trait individually (Conradsen et al., 2022), we observed the 12 population estimates of among-line (mutational) variance to vary in both magnitude and statistical support for all traits (Figure 2a,b; Table S1). There were no patterns of consistent change over time or divergence between the population size treatments (Figure 2a), suggesting the observed heterogeneity was not due to mutation-drift-selection processes. There was statistical support for 46 (63.9%) of the 72 estimates (6 traits in 12 population samples), with statistical support across all 12 samples for only one trait (ILD1.5, capturing proximo-posterior wing width: Figure 1b) (Figure 2a,b). Particularly for traits with fewer statistically supported estimates (Figure 2b) there were large differences in magnitude of among-line variance (ratio of largest to smallest ranged from 1.93 up to 34.4, median 7.6; Figure 2a). Nonetheless, there was little statistical support for among-line variance to differ between the population samples - estimates typically fell well within each other’s 90% CI, with only a few exceptions for traits ILD1.2 (two largest versus five smallest estimates) and ILD2.8 (largest versus smallest) (Figure 2a).

**Figure 2.**
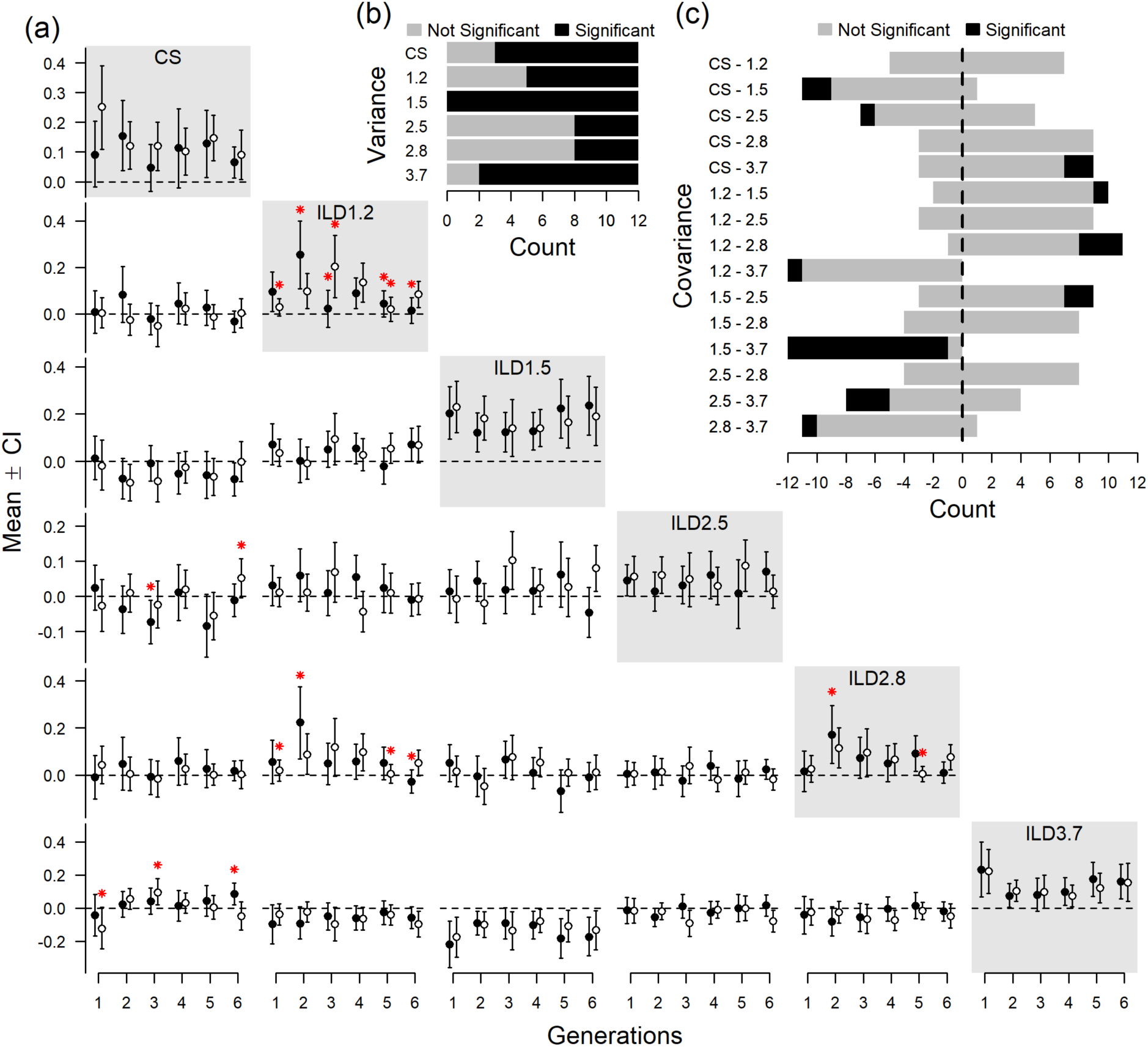
The variability among the 12 population sample estimates of among-line variance and covariance. (a) The REML estimate (point; REML-MVN bootstrap confidence intervals whiskers) of among-line variance (on diagonal, grey boxes; 90% CI) and covariances (below the diagonal; 95% CI) in each population size treatment (Small or Large, closed and open symbols, respectively) in each of six generations. The horizontal dashed line indicates zero; statistical support was concluded when CIs did not include zero (a one-tailed test for variances and two-tailed for covariances). Red stars highlight estimates where CI did not overlap with at least one other estimate of the same parameter. Inset plots are the counts of the 12 estimates of (b) variance for each trait or (c) pairwise covariances between traits that were (black) or were not (grey) statistically supported as different from zero. For the covariances, the counts of estimates are further plotted by sign (negative or positive). See Figure 1b for trait definitions and Table S1 for the parameter estimates. NB: two estimates of covariance between ILD2.5 and ILD3.7 were exactly 0; these were classified as positive.

There was considerably less statistical support for mutational covariance among traits, with only 15% (27 of 180) of pairwise covariances statistically distinct from zero (Figure 2a,c). Nonetheless, the average absolute correlation was 0.51 (Table S1). This pattern of frequent strong association between traits suggests that the relatively large sampling error of covariance estimates, rather than a lack of correlated mutational effects among traits, might contribute to the limited statistical evidence for mutational covariance. A binomial test of the null hypothesis of an equal number of positive and negative values rejects the null hypothesis (*P* = 0.0193) when 10 of 12 estimates have a consistent sign; six (of 15, 40%) mutational correlations were thus statistically supported as non-zero (Figure 2a,c) across the total dataset. There was, however, marked heterogeneity among the 12 repeated estimates of each pairwise covariance (correlation). The correlation estimates spanned the full range (−1 to 1) for one trait pair (ILD1.2: ILD2.8), with non-overlapping 95% CI of estimates for this and two other trait pairs (CS: ILD1.2 and CS: ILD3.7) (Figure 2a,c; Table S1).

### Common subspace and major axis of mutational variation

The distribution of eigenvalues of each population sample’s among-line covariance matrix was consistent with mutation generating trait covariance, and with variation in mutational effects among traits. Specifically, the large (relative to variance of individual traits) first eigenvalue (Figure 3; Table 1 vs Table S1) suggests traits share variation, while the robust statistical support for two axes of mutational variance in most population samples (Table S2) suggested that not all mutations generated the same correlated effects across traits. Here, we consider whether the similar pattern of eigenvalues (Figure 3) and low (relative to individual traits) heterogeneity among the 12 replicate samples in their eigenvalues (ratio of largest to smallest population sample estimate: 1.92, 2.90 and 3.08 for *m*_1_, *m*_2_ and *m*_3_, respectively: Table 1) also reflect concordance of the associated eigenvectors.

**Figure 3.**
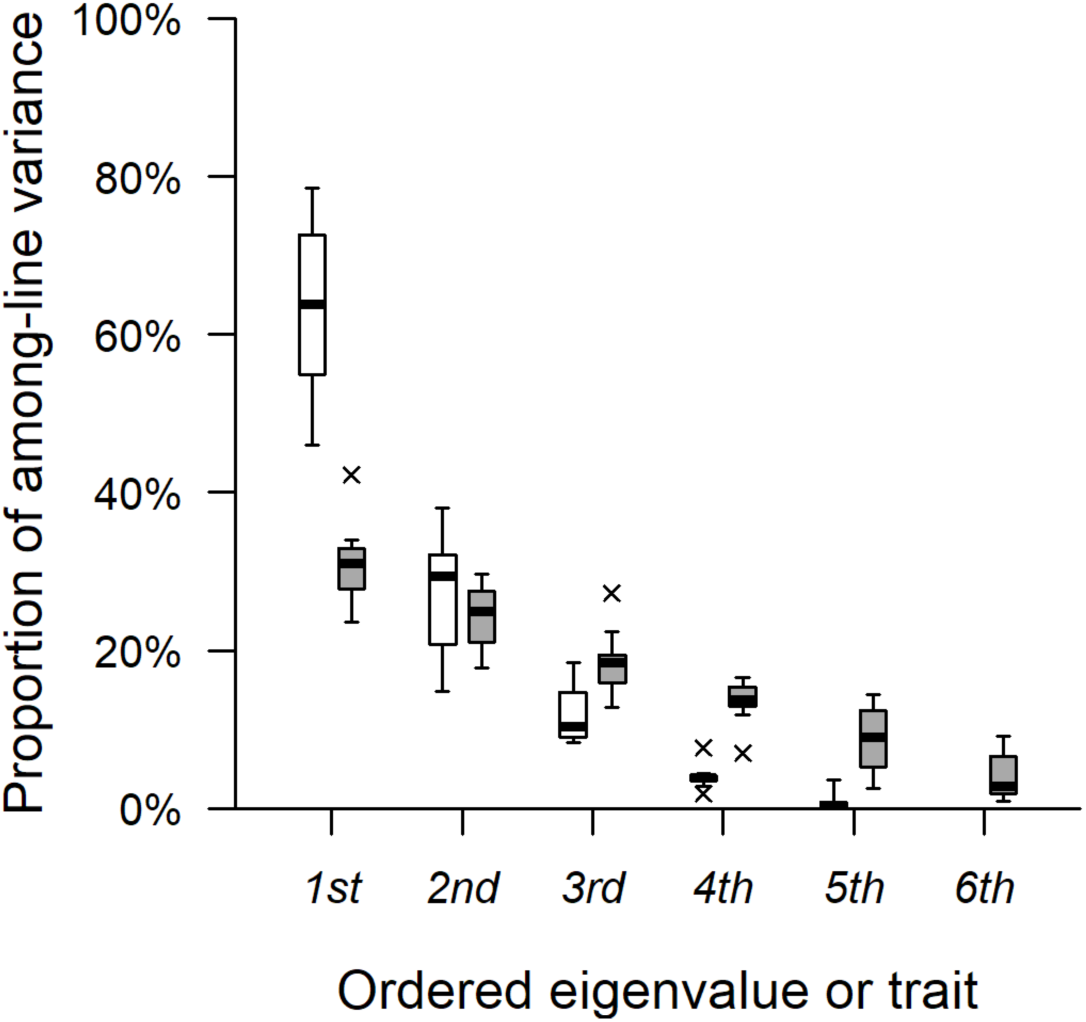
The distribution of among-line variation of six wing traits in 12 population samples. Open boxplots represent distribution across the 12 **M** of the ranked eigenvalues (largest to smallest, x-axis 1^st^ to 6^th^), plotted as the proportion of the variance in their respective **M**. For comparison, among-line variances of individual traits (diagonals of Table S1 and Figure 2a) were ranked from largest to smallest within each population sample, and their distribution also plotted (grey boxplots). Note, the combination of the six traits (i.e., eigenvector) associated with a specific ordered eigenvalue (1^st^ through 6^th^), and the specific trait ranked 1^st^ through 6^th^ may differ among the 12 matrices. For each boxplot, the median of the distribution is shown as a black band, boxes represent the interquartile range, whiskers represent 1.5 times the interquartile range, and estimates that fell outside this range are designated with crosses.

The eigenvalue of the major axis (*h*_1_) of the Krzanowski common subspace was within the confidence bounds estimated from the null datasets simulating a single population, and close to the maximum value (12) observed when the vector is present in the sampled subspaces of all 12 populations (Table 2). Consistent with this interpretation that each of the 12 populations had estimated the same multivariate axis of mutational input, there was statistically supported variance along *h*_1_ in all 12 population samples (Figure 4a). This common axis of wing shape variation, representing the best estimate of the multivariate wing shape most affected by mutation in the experiment, reflected an axis of expansion versus contraction of the proximal versus distal wing length (strongest, opposing, loadings on *h*_1_ from ILD1.5 and ILD3.7: Table 2). Notably, these two traits had the most consistent statistical support for, and least heterogeneity in magnitude among, among-line variance estimates, and were significantly negatively correlated in all but one population sample (average bounded correlation coefficient = −0.87: Figure 2; Table S1).

**Figure 4.**
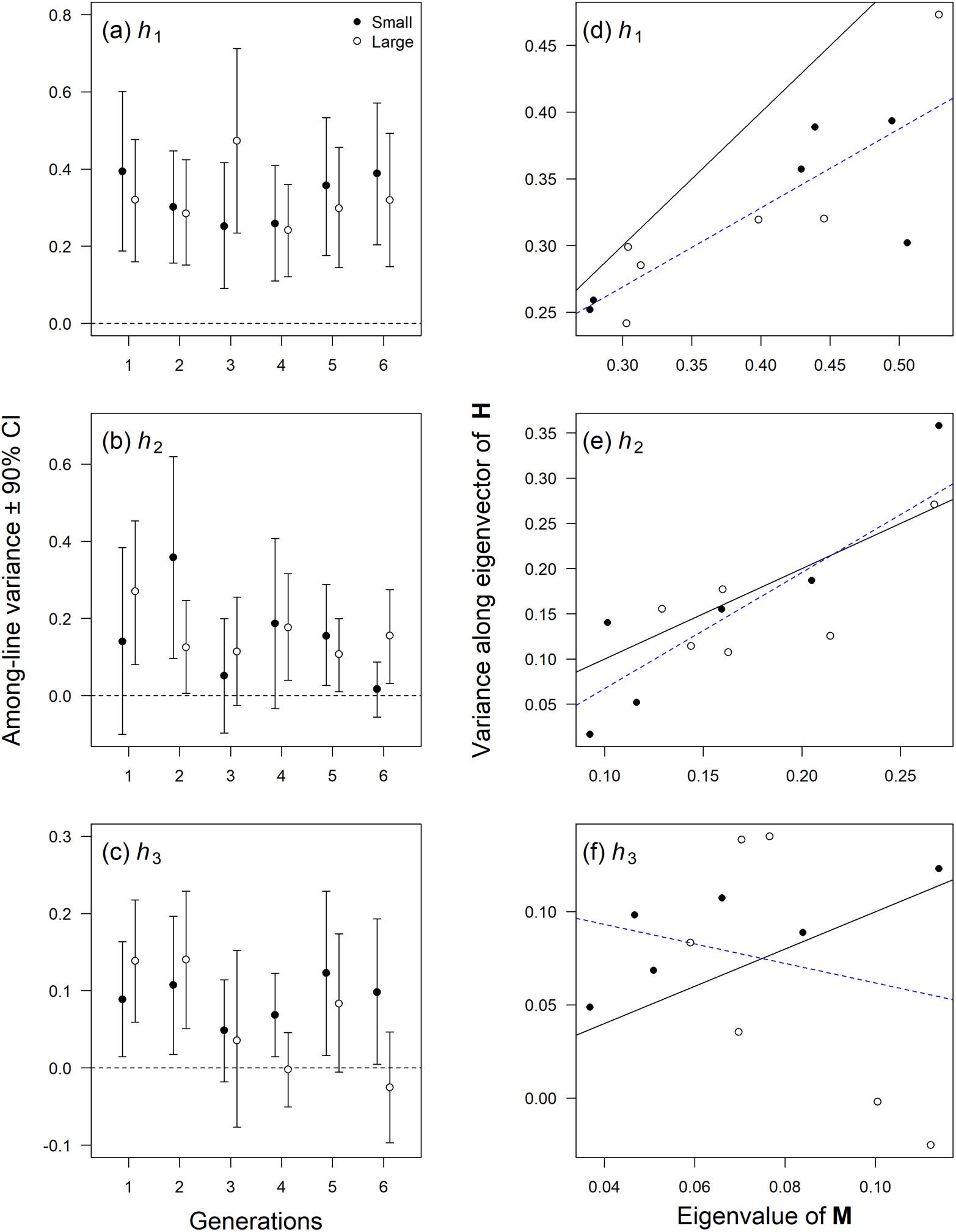
The among-line variance associated with each of the three eigenvectors of the Krzanowski’s subspace. Left-hand panels: The magnitude of the among-line variance (90% REML-MVN CI) is plotted for (a) *h_1_*, (b) *h_2_* and (c) *h_3_*. Within each plot, the populations are ordered by generation (x-axis), then by population size treatments (Small or Large, closed and open symbols, respectively). The dashed lines indicate zero. Note that the y-axis range differs among panels, reflecting decreasing total variance associated with each successive common subspace vector (a to c). Right-hand panels: the magnitude of variance along eigenvectors of **H** are plotted against the same-rank eigenvalue of each population sample’s respective **M**. (a) *h_1_* vs *m_1_*, (b) *h_2_* vs *m_2_* and (c) *h_3_* vs *m_3_*. Population size treatment is again indicated by symbol (Small or Large, closed and open symbols, respectively). The dashed blue line indicates the regression slope; solid grey line is the 1:1 line indicating equal magnitude along x and y axes.

**Table 2.** Eigenvalues and eigenvectors of the Krzanowski subspace matrix, H. The 95% CI (below estimate)**\*** for the eigenvalues (*λ*) of the observed data **H** were determined from Krzanowski subspace analyses applied to each of the 10,000 REML-MVN samples of each of the 12 **M**. For the null data **H**, the reported eigenvalue is the mean (and 95% CI are the 5% quantiles) of the Krzanowski subspace analyses applied to the 12 **M** estimated from the 1000 randomised datasets.

| | $h_1$ | $h_2$ | $h_3$ |
| --- | --- | --- | --- |
| $\lambda$ | 11.62<br>10.35; 11.49 | 8.86<br>6.87; 9.27 | 6.62<br>5.16; 7.14 |
| Null $\lambda$ | 11.38<br>10.61; 11.79 | 10.93<br>9.35; 11.66 | 8.56<br>5.77; 10.88 |
| CS | -0.37 | 0.76 | -0.32 |
| ILD1.2 | 0.27 | 0.41 | 0.36 |
| ILD1.5 | 0.65 | -0.05 | -0.46 |
| ILD2.5 | 0.20 | -0.03 | 0.64 |
| ILD2.8 | 0.16 | 0.42 | 0.34 |
| ILD3.7 | -0.55 | -0.27 | 0.19 |

\* Eigenvalues of $\mathbf{H}$ are bounded statistics (ranging from 0 to $p$ – here, 0 to 12). In conjunction with the nature of sampling error, which independently affects each estimated $\mathbf{M}$ and thus contributes dissimilarity, this has resulted in the eigenvalue of $h_1$ falling above its upper REML-MVN CI. Importantly, the observed $\lambda_1$ is within the range observed for the null $\lambda_1$ , which estimates the error in $h_1$ when the 12 samples of $\mathbf{M}$ are all drawn from the same population.

The major axis of multivariate mutational input, *h*_1_, was typically captured by the leading eigenvector of a population sample’s **M** (i.e., was aligned with *m*_1_). This is evidenced by: i) *h*_1_ being most closely aligned to *m*_1_ in 10 of the 12 populations (Table S4); ii) the magnitude of among-line variance being greater along *h*_1_ than along *h*_2_ or *h*_3_ in 11 populations (Figure 4a,b) and; iii) *h*_1_ typically accounting for almost as much variance in a population’s covariance matrix as *m*_1_ did (median = 86% of *m*_1_: Figure 4d). This reduced variance along *h*_1_ relative to each *m*_1_ is expected as variance along *m*_1_ will be inflated by estimation error (Blows & McGuigan, 2015), unique to that specific population sample. In population S2, *h*_1_ was more closely aligned with *m*_2_ (Table S4), reflected in less variance along *h*_1_ than *h*_2_ (0.30 versus 0.36: Figure 4a,b) and *h*_1_ accounting for only 60% as much variance as *m*_1_ (lower right point in Figure 4d). Nonetheless, even for this relatively incongruent population sample, *m*_1_ did capture a substantial signal of the major axis of mutational variance (*h*_1_) (Figure 4a,d). Similar to the observation of reduced heterogeneity (relative to individual traits) among eigenvalues of **M**, the largest estimate of among-line variance along *h*_1_ was 1.96 times greater than the smallest estimate, with estimates typically well within the CI of all other population samples (Figure 4a).

The eigenvalues (8.9 and 6.6: Table 2) of the other two interpretable common subspace axes implicated further axes of mutational input, with non-zero variance along these axes in most populations (Figure 4b,c). However, the absence of statistically supported variance along these axes in some population samples suggests that robust estimation of this variation may be limited by factors such as estimation error (increased relative to *h*_1_ due to the smaller magnitude of effect), segregating mutations, or micro-environmental heterogeneity. There were no clear temporal or among population size treatment patterns in the magnitude of variance along *h*_2_ or *h*_3_ (Figure 4b,c; Table S4). Notably, *h*_2_ primarily reflected mutational variance in wing size (CS), while *h*_3_ was most strongly influenced by ILD2.5 (Table 2): these are the only two traits for which individual trait analyses presented by Conradsen et al. (2022) robustly rejected the null hypothesis that among-line variance was homogenous among generations or population size treatments. ILD1.2 and ILD2.8 also contributed to *h*_2_ (larger versus smaller wings had disproportionally longer versus shorter anterior margins: Table 2, Figure 1b). Low concordance among the 12 estimates of variances and covariance involving these traits (Figure 2) suggest we have least confidence in the mutational effects on these traits, but shared signal (i.e., correlated pleiotropic effects) with other traits may underpin their contribution to this axis of wing shape variation, observed in most population samples (Figure 4b).

## DISCUSSION

Accurately estimating mutational effects, and particularly pleiotropic effects across traits, represents a clear challenge to understanding how factors such as genetic background, environmental context or the traits under consideration might each influence mutational characteristics (Conradsen et al., 2022). However, by leveraging shared signal against independent error, multivariate analyses of mutational covariance across traits may potentially provide more robust insights than interrogation of mutational correlations of individual trait pairs. Here, we provide support for this through re-analysis of published data from an experiment that was: i) designed to reflect typical levels of sampling error of a spontaneous mutation accumulation experiment (Conradsen et al., 2022) and; ii) is in a system (Drosophild wings) where shared mutational signal is expected (extensive developmental and genetic evidence of pleiotropic alleles and trait covariances: Carreira et al., 2011; Dugand et al., 2021; Houle et al., 2017; Matamoro-Vidal et al., 2015; Pitchers et al., 2019). Our results suggest that while mutational differences may be detectable for major axes of mutational variance, for other multivariate traits this may remain difficult. We discuss these points below, and highlight the value of replicated estimation for deepening our understanding of mutational characteristics.

Estimating mutational (co)variances repeatedly from independent samples of the same mutation accumulation experiment allowed us to confidently conclude that there was a strong signal of mutation generating trait covariance. Notably, although all 12 matrices captured variation along the same axis of wing shape (*h*_1_: Table 2; Figure 4a), the trait covariance contributing most strongly to this robustly estimated axis of mutational variance was not unequivocally identifiable *a priori* from a single population sample. We expected the magnitude of mutational (co)variance would predict the extent of concordance across the 12 estimates. However, the covariance contributing most strongly to *h*_1_ (ILD1.5: ILD3.7), while the most consistently statistically supported across samples (Figure 2c), was the strongest of the statistically supported trait correlations in only one population sample. Similarly, while the two traits individually had relatively large average estimates of among-line variance, other traits had the largest (statistically supported) estimate in some samples (Figure 2; Tables S1) (also see Figure 7 versus Figure 5 of Conradsen et al., 2022). Thus, the magnitude of estimated (co)variance in any single sample of mutational phenotypes may not provide a simple diagnostic of the strongest mutational signal.

Among-line variation along *h*_1_ was predominantly captured by variation along the major axis of the among-line covariance matrix estimated within each population sample (i.e., *m*_1_). Higher variance along population-specific *m*_1_ than along *h*_1_ (Figure 4d) is consistent with the Krzanowski common subspace analysis finding the shared signal of mutational input, with independent sampling error within each population sample leading to overestimation of the magnitude of mutational variance associated with the most mutationally variable multivariate trait. Leading eigenvalues of genetic (and mutational) covariance matrices are particularly vulnerable to inflation by sampling error (Blows & McGuigan, 2015; Hill & Thompson, 1978; McGuigan et al., 2014). Statistical approaches can estimate the inflation on eigenvalues (McGuigan et al., 2014), although the empirical approach of repeated estimation from independent samples applied here will provide greater certainty as to the orientation of the eigenvectors.

Here, concordant patterns across replicate independent samples of the same mutation accumulation experiment provided considerable confidence that these patterns reflect the underlying mutational signal. We suggest that concordance of mutational covariances (or major axes of mutational variance) across independent samples of the mutational spectrum (e.g., independent mutation accumulation experiments) may help distinguish the contribution of pleiotropy from that of linkage, leading to deeper insight into general characteristics of the distribution of pleiotropic effects. In mutation accumulation experiments, all mutations arising within a line are in linkage. Particularly for traits with directionally biased mutational effects (e.g., fitness traits, which evolve lower values under mutation-drift: Halligan & Keightley, 2009), linkage of mutations with independent, but directionally biased, effects on each trait may contribute substantially to among-line covariances (Estes et al., 2005). For traits without biased directional effects, including *Drosophila* wing traits (Houle & Fierst, 2013; Santiago et al., 1992), strong trait covariances may still arise via linkage of randomly sampled mutations. However, chance linkage of randomly sampled mutations is not expected to cause consistent covariance patterns across independent mutation accumulation experiments. Rather, in the absence of directional bias to the distribution of mutational effects, the average mutational covariance generated by linkage should approach zero. Repeated observation of the same major axes of phenotypic variation in independent samples of mutation therefore suggest conserved patterns of correlated pleiotropic effects of mutations.

The major axes of mutational input to wing shape estimated in the current study (*h*_1_, Table 2) suggests that mutation predominantly generated variation along an axis of shorter and wider versus longer and narrower wing tips. Intriguingly, this was qualitatively the same mutational wing shape variance identified in independent studies of *Drosophila* wings, using different experimental designs and analytical approaches. Dugand et al. (2021) estimated mutational (co)variance in an outbred, randomly mating, *D. serrata* population founded by the DsGRP lines (including the ancestor line of the mutation accumulation lines considered in the current study). In that outbred population analysis, *m*_1_ was characterized by lengthening ILD1.5 and ILD5.7 while shortening along ILD4.6 and ILD3.9 (and vice versa: see their Table 3). Similarly, Houle et al. (2017) estimated the major axis of mutational variance using inbred mutation accumulation lines of *D. melanogaster* (see their Extended Data Figure 1), finding that mutation pushed out the landmarks corresponding to LM5 and LM7 in the current study (increasing ILD1.5 and ILD1.7), while pulling closer together landmarks corresponding to LM3,4 and LM8,9 (shortening distances between those landmarks, including ILD3.7, ILD3.9 and ILD4.6).

The apparent consistency of multivariate mutational covariance patterns across these different studies of *Drosophila* wings suggests bias in either the frequency or effect sizes of pleiotropic mutations, driving persistent bias in the phenotypic variation generated by new mutation. This inference of conserved mutational effects is qualitative rather than quantitative. Conservation of mutational effects among taxa is also implicated by evidence that the mutational covariance matrix estimated in one species (*D. melanogaster*) strongly predicted the pattern of divergence among 112 Drosophilidae species over 40 million years (Houle et al., 2020; Houle et al., 2017). In contrast, Houle and Fierst (2013) compared **M** estimated from two genotypes of *D. melanogaster* and concluded that these **M** were quite different from one another, including in the orientation of *m*_1_. Few other studies have quantitatively compared **M** estimated from independent samples of mutations. François Mallard et al. (2023) estimated **M** of locomotor behavior traits in two genotypes of *Caenorhabditis elegans*, finding no evidence that these independently sampled **M** diverged.

Direct, quantitative, comparison of **M** from different studies is hampered by methodological differences (here, for example, differences in wing landmarks visible under different imaging methods). Inconsistent statistical support for mutational variance of individual traits can also limit direct comparability. Although Conradsen et al. (2022) measured the same traits as Dugand et al. (2021), differences in which traits had non-zero mutational variance estimates, coupled with the impact of zero variance on multivariate model convergence, led to different traits being included in analyses of **M**. Whether inconsistency among independent studies in the detection of mutational variance for individual traits reveals genotypic differences in mutational effects will require further comparative data to assess the role of sampling error.

Even when mutational variance is observed for individual traits, mutational variance co-variance matrices are characterised by so-called nearly null spaces (Gomulkiewicz & Houle, 2009) – that is, **M** typically have one or more very small eigenvalues, statistically indistinguishable from zero (Dugand et al., 2021; Estes et al., 2005; Hine et al., 2018; Houle & Fierst, 2013; Latimer et al., 2014; François Mallard et al., 2023; Miller et al., 2023). While this evidence that mutational input to phenotypic variation is uneven across phenotypic trait space is interesting on its own, it is challenging to investigate further the traits with low mutational input. Comparative approaches may again extend our understanding of this question. Houle and Fierst (2013) found that, despite differences between *D. melanogaster* genotypes in the multivariate traits with high mutational input, the traits associated with low mutation were concordant. Dugand et al. (2021) found that *D. serrata* wing phenotypes with low (undetectable) mutational input also have very low standing genetic variance. These observations suggest that it is not simply chance sampling of mutation or estimation error that causes low mutational input to some multivariate traits, and opens the path to further investigate why such variation might be relatively inaccessible.

Overall, our results suggest that even relatively modest mutation accumulation experiments may provide valuable information on the multivariate axes of traits space to which mutation contributes the greatest new variance each generation. Such insights are valuable, particularly for deepening our understanding of how mutation and selection interact to shape standing genetic variance, and long-term responses to selection – for example, does mutation predominantly introduce variance to traits with relatively high standing genetic variation (as observed for Drosophildae wings: Dugand et al., 2021; Houle et al., 2017) or is there a mismatch between mutational and standing genetic variance implicating selection acting against new mutations (for example, as observed for thermal performance traits: Latimer et al., 2014; or locomotor behaviour traits: F Mallard et al., 2023). Replication of estimates both within the same experiment and across mutation accumulation experiments are a powerful approach to both identify dominant patterns of pleiotropic correlations, and to ensure robust inference of how that distribution of effects may vary.

## DATA AVAILABILITY

Data analysed in this paper were published by (Conradsen et al., 2021)

## ACKNOWLEDGMENTS

The authors thank Stephen F. Chenoweth for providing us with the *D. serrata* MA lines, and Adam Reddiex, Nicholas Appleton, Jack Price and Derek Sun for their help with maintaining the flies and collecting the data. The authors also thank Emma Hine for helpful discussions about analyses.

## CONFLICTS OF INTEREST

None declared.

## SUPPLEMENTARY MATERIALS

**Table S1.**
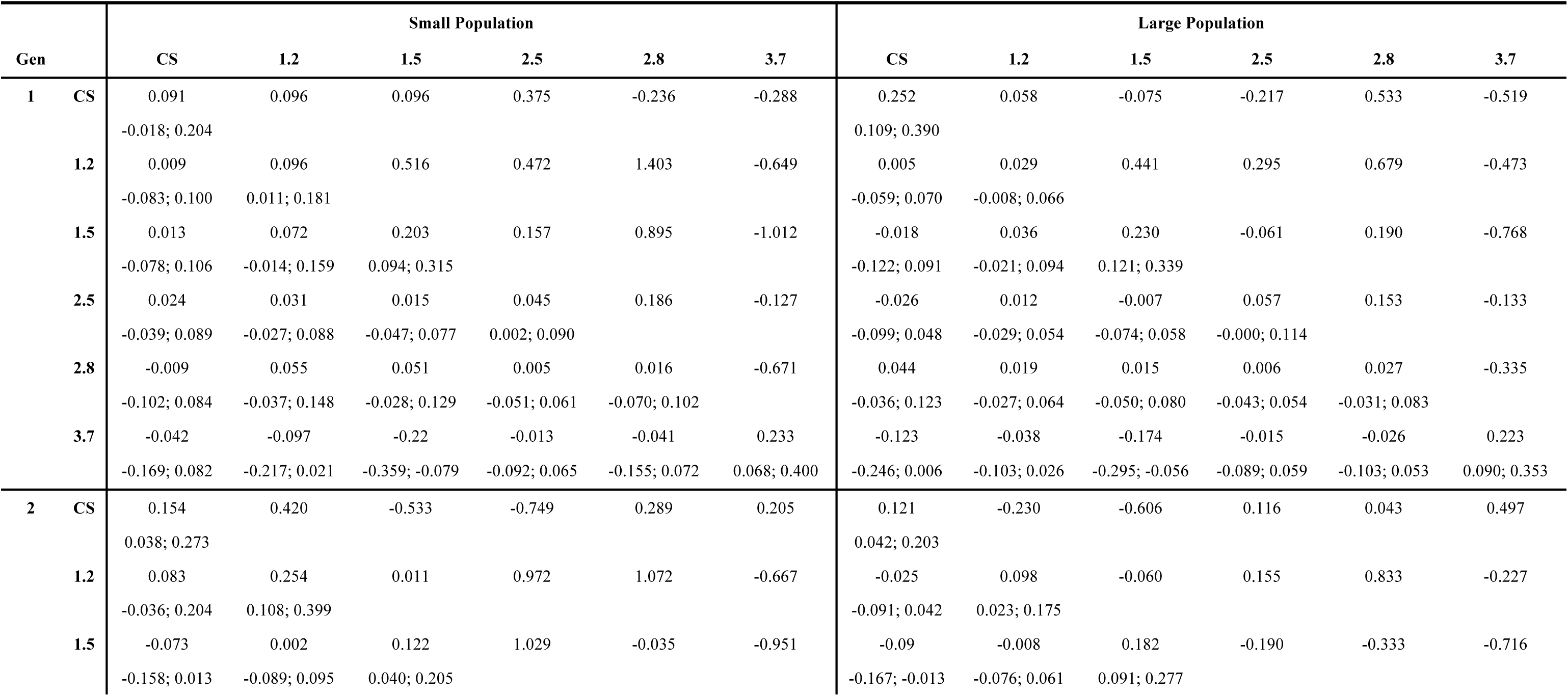
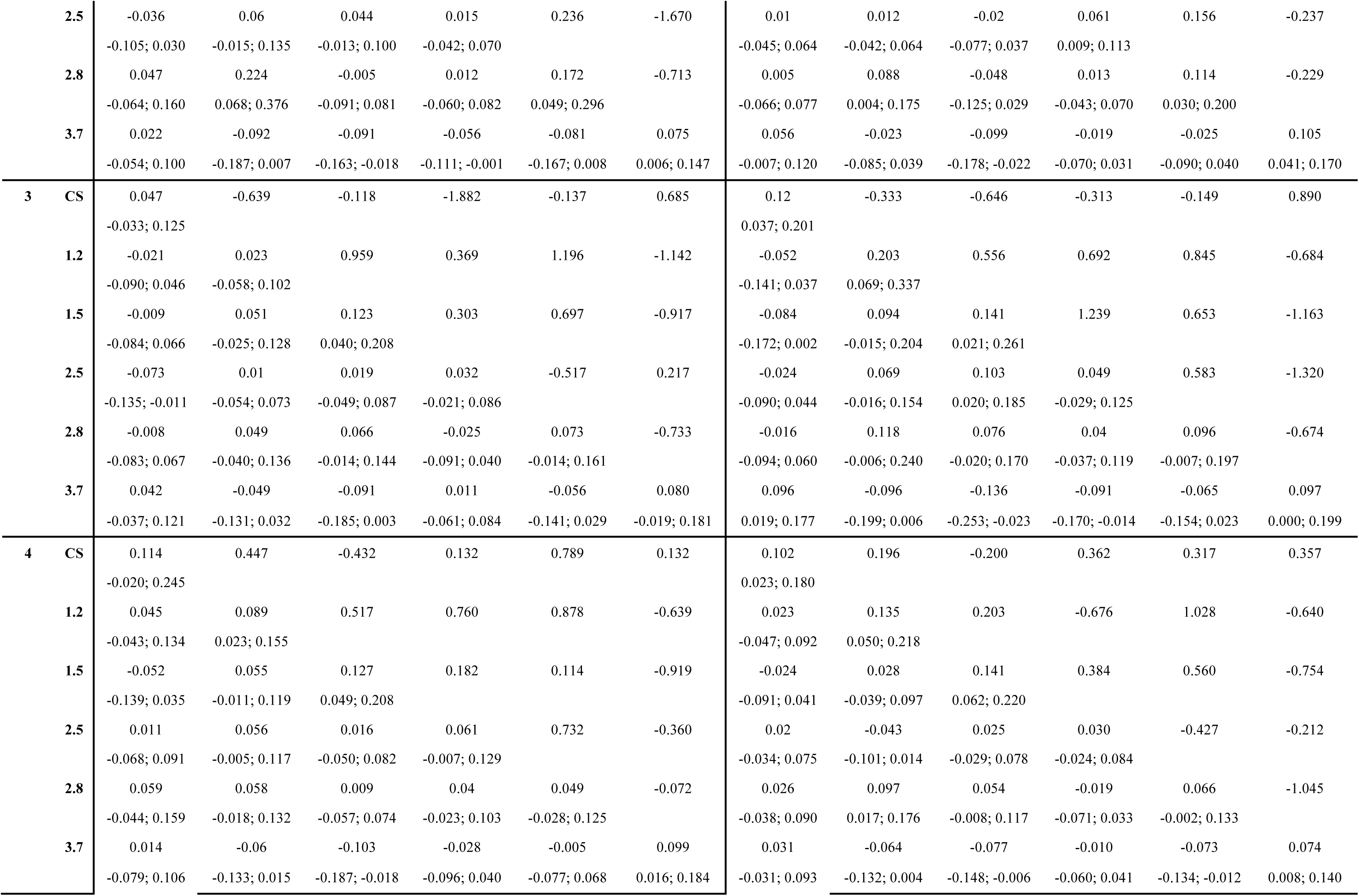
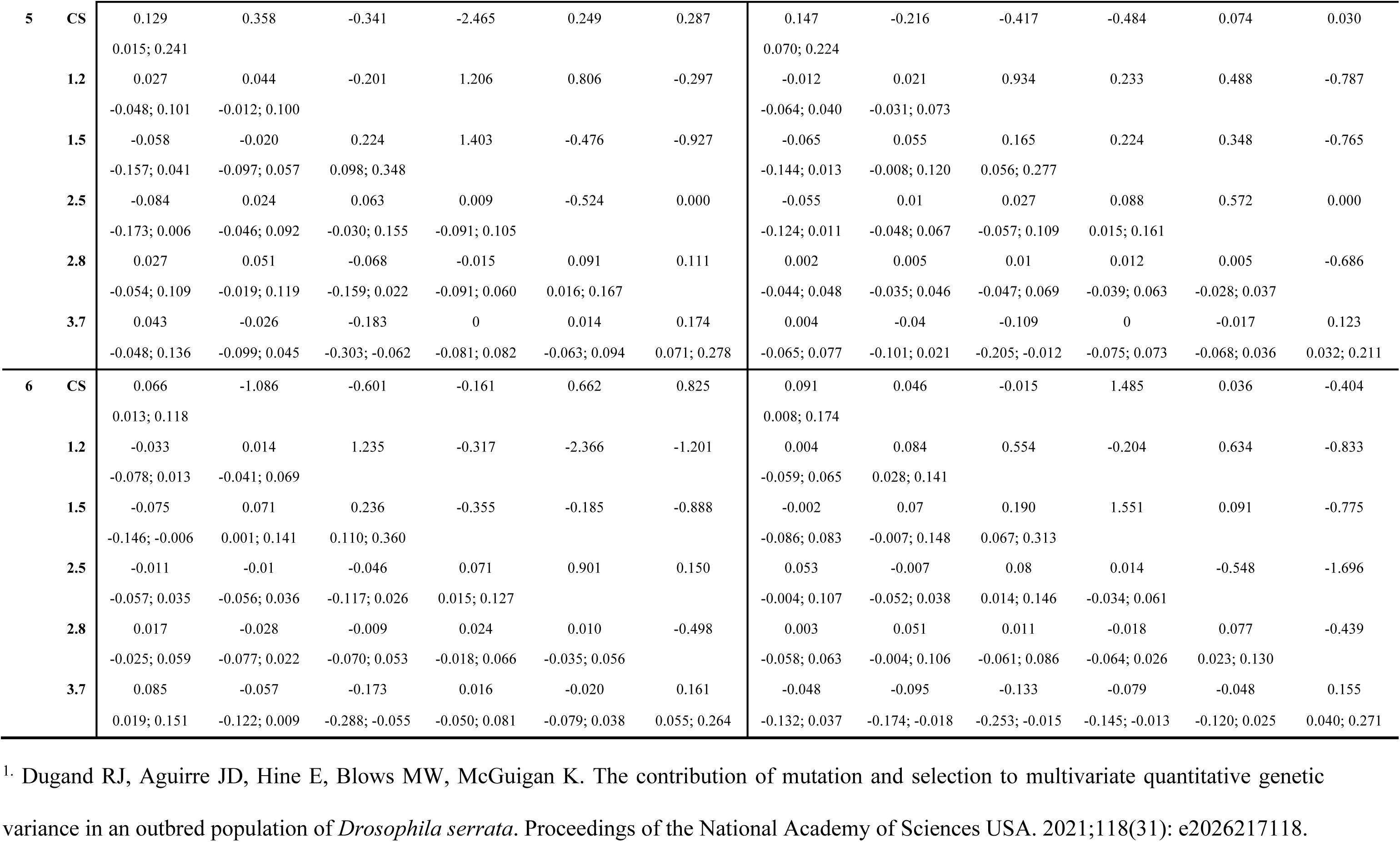
Mutational variances (diagonal), covariances (lower off-diagonal) and correlations (upper off-diagonal) for the small and large population treatments, for each of the six generations, for six wing traits. Confidence intervals (below variance and covariance estimates) were determined using REML-MVN, and are 90% for variances (for which negative values are not possible: the frequency of negative REML-MNV estimates equate to the LRT probability^1^) and 95% for covariances (reflecting a two-tailed test for a parameter that can be positive or negative). No confidence intervals are reported for correlations because some MVN samples of variances are negative, precluding calculation. To calculate the average absolute strength of correlations, estimates exceeding theoretical bounds (−1 and 1) were set to |1|.

**Table S2.** Factor analytic modelling of among-line covariance in each of the 12 populations. Sequential REML mixed models (model 1) were fit with zero up to six dimensions of among-line variance (columns, left to right; the number of among-line parameters estimated by the model is shown in parentheses on the second row). The −2 loglikelihood (−2LL) of each model is reported. Statistical support for each additional dimension was determined by log-likelihood ratio tests (*P*-values shown below the −2LL for that dimension); the difference in −2LL between sequential models follows a chi-squared distribution with degrees of freedom determined by the difference in number of parameters. *P*-values are shown in bold where the null hypothesis (variance along that dimension is not greater than zero) was rejected.

| Treat | Gen | None | 1 | 2 | 3 | 4 | 5 | 6 |
| --- | --- | --- | --- | --- | --- | --- | --- | --- |
|  |  | (0) | (6) | (11) | (15) | (18) | (20) | (21) |
| Small | 1 | 6218.8 | 6194.3<br>**0.0004** | 6184.3<br>*0.0752* | 6176.4<br>*0.0953* | 6175.6<br>*0.8495* | 6175.6<br>*>0.9999* | 6175.6<br>*>0.9999* |
|  | 2 | 6474.0 | 6444.6<br>**0.0001** | 6430.7<br>**0.0163** | 6428.0<br>*0.6092* | 6427.1<br>*0.8254* | 6427.1<br>*>0.9999* | 6427.1<br>*>0.9999* |
|  | 3 | 5676.5 | 5660.3<br>**0.0127** | 5647.4<br>**0.0243** | 5646.7<br>*0.9513* | 5646.7<br>*>0.9999* | 5646.1<br>*0.7408* | 5646.2<br>*0.7518* |
|  | 4 | 6603.6 | 6588.6<br>**0.0203** | 6577.4<br>**0.0476** | 6575.6<br>*0.7725* | 6574.5<br>*0.7771* | 6574.5<br>*>0.9999* | 6574.5<br>*>0.9999* |
|  | 5 | 5489.5 | 5466.3<br>**0.0007** | 5460.4<br>*0.3161* | 5456.1<br>*0.3669* | 5455.9<br>*0.9776* | 5452.7<br>*0.2019* | 5452.7<br>*>0.9999* |
|  | 6 | 6633.6 | 6590.0<br>**<0.0001** | 6578.5<br>**0.0423** | 6573.9<br>*0.3309* | 6572.1<br>*0.6149* | 6572.1<br>*>0.9999* | 6572.1<br>*>0.9999* |
| **Large** | 1 | 6473.9 | 6448.4<br>**0.0003** | 6429.6<br>**0.0021** | 6420.4<br>*0.0563* | 6417.2<br>*0.3618* | 6416.5<br>*0.7047* | 6416.5<br>*>0.9999* |
|  | 2 | 6833.0 | 6795.8<br>**<0.0001** | 6785.8<br>*0.0752* | 6775.9<br>**0.0421\*** | 6771.0<br>*0.1793* | 6769.8<br>*0.5488* | 6769.8<br>*>0.9999* |
|  | 3 | 5498.8 | 5477.7<br>**0.0018** | 5457.5<br>**0.0011** | 5453.7<br>*0.4337* | 5452.9<br>*0.8495* | 5452.9<br>*>0.9999* | 5452.8<br>*0.7518* |
|  | 4 | 6415.9 | 6390.3<br>**0.0003** | 6377.7<br>**0.0274** | 6371.6<br>*0.1918* | 6371.5<br>*0.9918* | 6371.5<br>*>0.9999* | 6370.7<br>*0.3711* |
|  | 5 | 6047.7 | 6032.6<br>**0.0195** | 5998.9<br>**<0.0001** | 5994.9<br>*0.4060* | 5994.0<br>*0.8254* | 5994.0<br>*>0.9999* | 5994.0<br>*>0.9999* |
|  | 6 | 5455.0 | 5435.1<br>**0.0029** | 5424.6<br>*0.0622* | 5417.4<br>*0.1257* | 5413.2<br>*0.2407* | 5413.2<br>*>0.9999* | 5413.1<br>*0.7518* |

\* statistical support for three but not two dimensions suggests re-ordering such that the FA0(2) model estimated a lower variance subset of principal components (MEYER AND KIRKPATRICK 2008). For this population sample, we interpret two dimensions as being statistically supported.

Meyer, K., and M. Kirkpatrick. Perils of parsimony: Properties of reduced rank estimates of genetic covariance matrices. Genetics 2008; 180:
1153-1166.

**Table S3.** The eigenvectors for M. Eigenanalyses were performed separately for **M** in each population (small, S, or large, L for every generation). The corresponding eigenvalues are reported in Table 2.

| **Population** | **Trait** | $m_1$ | $m_2$ | $m_3$ | $m_4$ | $m_5$ | $m_6$ |
| --- | --- | --- | --- | --- | --- | --- | --- |
| **S1** | **size** | 0.10 | 0.90 | 0.14 | 0.26 | -0.28 | 0.06 |
|  | **1.2** | 0.31 | 0.01 | -0.77 | 0.33 | 0.16 | 0.42 |
|  | **1.5** | 0.63 | -0.20 | 0.26 | -0.31 | -0.47 | 0.43 |
|  | **2.5** | 0.07 | 0.34 | -0.40 | -0.82 | 0.11 | -0.18 |
|  | **2.8** | 0.16 | -0.16 | -0.34 | 0.17 | -0.61 | -0.65 |
|  | **3.7** | -0.68 | -0.05 | -0.21 | -0.14 | -0.53 | 0.42 |
| **S2** | **size** | 0.21 | 0.60 | 0.75 | 0.00 | -0.07 | -0.18 |
|  | **1.2** | 0.72 | 0.04 | -0.09 | -0.37 | 0.37 | 0.45 |
|  | **1.5** | 0.03 | -0.63 | 0.52 | 0.27 | 0.50 | -0.10 |
|  | **2.5** | 0.12 | -0.27 | 0.02 | -0.7 | -0.13 | -0.64 |
|  | **2.8** | 0.59 | 0.03 | -0.32 | 0.55 | -0.08 | -0.50 |
|  | **3.7** | -0.28 | 0.41 | -0.26 | -0.06 | 0.76 | -0.32 |
| **S3** | **size** | -0.18 | 0.68 | -0.34 | 0.08 | 0.21 | -0.58 |
|  | **1.2** | 0.33 | -0.03 | 0.22 | 0.22 | 0.89 | 0.08 |
|  | **1.5** | 0.63 | 0.06 | -0.7 | 0.16 | -0.12 | 0.27 |
|  | **2.5** | 0.05 | -0.67 | -0.18 | 0.32 | -0.02 | -0.65 |
|  | **2.8** | 0.43 | 0.31 | 0.53 | 0.51 | -0.39 | -0.16 |
|  | **3.7** | -0.53 | 0.00 | -0.16 | 0.74 | 0.02 | 0.37 |
| **S4** | **size** | -0.01 | 0.70 | 0.54 | -0.13 | 0.21 | 0.40 |
|  | **1.2** | 0.49 | 0.34 | -0.09 | 0.07 | -0.8 | -0.02 |
|  | **1.5** | 0.59 | -0.36 | 0.19 | 0.49 | 0.21 | 0.45 |
|  | **2.5** | 0.28 | 0.23 | -0.75 | -0.32 | 0.32 | 0.33 |
|  | **2.8** | 0.20 | 0.42 | -0.12 | 0.51 | 0.37 | -0.60 |
|  | **3.7** | -0.55 | 0.17 | -0.31 | 0.61 | -0.18 | 0.41 |
| **S5** | **size** | -0.29 | 0.64 | 0.52 | -0.32 | 0.12 | 0.34 |
|  | **1.2** | -0.04 | 0.35 | -0.44 | -0.55 | -0.49 | -0.37 |
|  | **1.5** | 0.72 | 0.10 | 0.18 | -0.33 | 0.48 | -0.32 |
|  | **2.5** | 0.17 | -0.30 | -0.32 | -0.49 | 0.07 | 0.73 |
|  | **2.8** | -0.21 | 0.39 | -0.62 | 0.21 | 0.61 | 0.02 |
|  | **3.7** | -0.57 | -0.46 | 0.11 | -0.44 | 0.37 | -0.34 |
| **S6** | **size** | -0.29 | 0.36 | 0.62 | 0.29 | -0.52 | 0.23 |
|  | **1.2** | 0.22 | 0.05 | -0.35 | 0.16 | -0.68 | -0.58 |
|  | **1.5** | 0.71 | 0.23 | 0.19 | 0.57 | 0.29 | 0.00 |
|  | **2.5** | -0.11 | -0.81 | 0.00 | 0.53 | -0.12 | 0.17 |
|  | **2.8** | -0.02 | -0.28 | 0.64 | -0.17 | 0.18 | -0.67 |
|  | **3.7** | -0.59 | 0.28 | -0.24 | 0.51 | 0.37 | -0.35 |

Table S3 (cont.)
| Population | Trait | $m_1$ | $m_2$ | $m_3$ | $m_4$ | $m_5$ | $m_6$ |
| --- | --- | --- | --- | --- | --- | --- | --- |
| **L1** | **size** | 0.42 | 0.81 | 0.09 | 0.02 | -0.09 | 0.39 |
|  | **1.2** | 0.12 | -0.07 | -0.28 | 0.60 | 0.71 | 0.21 |
|  | **1.5** | 0.55 | -0.57 | 0.28 | 0.13 | -0.29 | 0.43 |
|  | **2.5** | -0.01 | -0.08 | -0.87 | -0.19 | -0.28 | 0.33 |
|  | **2.8** | 0.11 | 0.10 | -0.17 | 0.69 | -0.52 | -0.46 |
|  | **3.7** | -0.70 | 0.04 | 0.21 | 0.34 | -0.24 | 0.54 |
| **L2** | **size** | -0.49 | 0.13 | -0.52 | 0.64 | 0.25 | -0.07 |
|  | **1.2** | 0.03 | -0.63 | 0.22 | 0.04 | 0.62 | -0.41 |
|  | **1.5** | 0.73 | 0.14 | -0.08 | 0.35 | 0.39 | 0.41 |
|  | **2.5** | -0.05 | -0.15 | -0.66 | -0.63 | 0.29 | 0.26 |
|  | **2.8** | -0.12 | -0.70 | 0.00 | 0.24 | -0.33 | 0.57 |
|  | **3.7** | -0.46 | 0.25 | 0.49 | -0.12 | 0.45 | 0.52 |
| **L3** | **size** | -0.3 | 0.54 | -0.69 | 0.15 | 0.01 | -0.34 |
|  | **1.2** | 0.51 | 0.60 | 0.39 | 0.45 | 0.15 | -0.06 |
|  | **1.5** | 0.49 | -0.28 | -0.40 | -0.09 | 0.71 | -0.12 |
|  | **2.5** | 0.31 | -0.08 | -0.46 | 0.36 | -0.35 | 0.67 |
|  | **2.8** | 0.33 | 0.43 | -0.08 | -0.80 | -0.17 | 0.18 |
|  | **3.7** | -0.45 | 0.27 | 0.10 | -0.04 | 0.57 | 0.62 |
| **L4** | **size** | -0.01 | 0.55 | 0.76 | -0.05 | -0.11 | 0.34 |
|  | **1.2** | 0.56 | 0.48 | -0.24 | -0.13 | 0.62 | -0.04 |
|  | **1.5** | 0.48 | -0.58 | 0.37 | -0.55 | 0.05 | 0.06 |
|  | **2.5** | -0.06 | -0.23 | 0.48 | 0.51 | 0.5 | -0.45 |
|  | **2.8** | 0.49 | 0.20 | 0.06 | 0.13 | -0.58 | -0.6 |
|  | **3.7** | -0.47 | 0.21 | 0.07 | -0.64 | 0.12 | -0.55 |
| **L5** | **size** | -0.39 | 0.72 | -0.45 | -0.32 | 0.11 | -0.04 |
|  | **1.2** | 0.23 | 0.11 | -0.07 | -0.23 | -0.92 | -0.15 |
|  | **1.5** | 0.71 | 0.16 | 0.11 | -0.59 | 0.31 | 0.13 |
|  | **2.5** | 0.20 | -0.45 | -0.84 | -0.03 | 0.10 | -0.18 |
|  | **2.8** | 0.06 | 0.04 | -0.23 | 0.17 | -0.16 | 0.94 |
|  | **3.7** | -0.49 | -0.48 | 0.11 | -0.69 | -0.05 | 0.19 |
| **L6** | **size** | 0.14 | 0.48 | 0.70 | 0.00 | 0.46 | 0.22 |
|  | **1.2** | 0.35 | -0.49 | 0.13 | -0.63 | 0.35 | -0.32 |
|  | **1.5** | 0.62 | 0.13 | -0.56 | 0.26 | 0.42 | 0.20 |
|  | **2.5** | 0.26 | 0.43 | 0.02 | 0.14 | -0.17 | -0.84 |
|  | **2.8** | 0.16 | -0.58 | 0.36 | 0.7 | 0.06 | -0.13 |
|  | **3.7** | -0.62 | 0.00 | -0.22 | 0.14 | 0.67 | -0.31 |

**Table S4.** Pairwise dot products between eigenvectors of M and the eigenvectors of H. The largest dot product (indicating the strongest relationship) is bolded for each population sample’s comparison for each **H** vector. Dot products of normalised vectors range in magnitude from 0 (orthogonal orientation) to 1 (identical orientation). Dot products can be positive or negative, but we focus here on the absolute magnitude for two reasons. First, eigenvector loadings have arbitrary direction from the origin along the defined axis such that the sign of all loadings can be flipped (positive to negative and negative to positive) with no effect on the interpretation of trait covariation. Second, eigenvectors of **H** are not constrained to be orthogonal within the space of each **M**, which can result in opposing sign of association with two orthogonal eigenvectors of **M** if the **H** vector falls between them. The eigenvectors in Table S2 were flipped such that the focal dot products (i.e., between same rank **H** and **M** eigenvectors) were typically positive. Note that no *m*_5_ or *m*_6_ eigenvectors from any population sample were included in the common subspace analysis, and *m*_4_ was included for only one matrix (Table 1). Dot products (*r*) can be converted to angle degree by: (𝑐𝑜𝑠^-1^𝑟)(^180^/_𝜋_).

| Population | $\mathbf{H}$<br>vector | $m_1$ | $m_2$ | $m_3$ | $m_4$ | $m_5$ | $m_6$ |
| --- | --- | --- | --- | --- | --- | --- | --- |
| S1 | $h_1$ | **0.87** | -0.39 | -0.11 | -0.27 | 0.06 | 0.00 |
| S2 | $h_1$ | 0.41 | **-0.90** | 0.13 | 0.06 | -0.01 | 0.09 |
| S3 | $h_1$ | **0.94** | -0.31 | -0.13 | -0.13 | 0.01 | 0.05 |
| S4 | $h_1$ | **0.91** | -0.38 | -0.10 | 0.07 | 0.07 | -0.12 |
| S5 | $h_1$ | **0.88** | 0.18 | -0.42 | -0.07 | 0.04 | -0.10 |
| S6 | $h_1$ | **0.93** | -0.33 | 0.03 | 0.10 | 0.00 | -0.12 |
| L1 | $h_1$ | 0.64 | **-0.71** | -0.24 | 0.12 | 0.03 | -0.11 |
| L2 | $h_1$ | **0.89** | -0.41 | -0.20 | -0.02 | 0.09 | 0.04 |
| L3 | $h_1$ | **0.93** | -0.32 | -0.06 | -0.03 | 0.09 | -0.15 |
| L4 | $h_1$ | **0.79** | -0.58 | -0.04 | 0.10 | 0.18 | 0.02 |
| L5 | $h_1$ | **0.99** | 0.05 | -0.05 | 0.07 | -0.07 | 0.07 |
| L6 | $h_1$ | **0.86** | -0.23 | -0.41 | 0.06 | -0.20 | -0.06 |
| S1 | $h_2$ | 0.42 | **0.63** | -0.30 | 0.48 | -0.24 | -0.18 |
| S2 | $h_2$ | **0.77** | 0.41 | 0.44 | 0.10 | -0.16 | -0.05 |
| S3 | $h_2$ | 0.29 | **0.65** | 0.14 | 0.15 | 0.36 | -0.57 |
| S4 | $h_2$ | 0.39 | **0.81** | 0.42 | -0.04 | 0.02 | -0.10 |
| S5 | $h_2$ | -0.21 | **0.92** | -0.07 | -0.23 | 0.02 | 0.20 |
| S6 | $h_2$ | -0.01 | 0.11 | 0.65 | 0.03 | **-0.71** | -0.26 |
| L1 | $h_2$ | 0.58 | **0.65** | -0.16 | 0.46 | 0.09 | 0.01 |
| L2 | $h_2$ | -0.32 | -0.52 | -0.41 | **0.64** | 0.16 | -0.15 |
| L3 | $h_2$ | 0.21 | **0.78** | -0.39 | -0.03 | -0.18 | -0.39 |
| L4 | $h_2$ | 0.53 | **0.68** | 0.45 | 0.15 | -0.12 | 0.15 |
| L5 | $h_2$ | -0.09 | **0.74** | -0.48 | -0.05 | -0.37 | 0.25 |
| L6 | $h_2$ | 0.45 | -0.10 | **0.82** | -0.02 | 0.32 | 0.08 |
| S1 | $h_3$ | 0.24 | 0.04 | **0.85** | 0.32 | -0.13 | 0.32 |
| S2 | $h_3$ | -0.40 | -0.03 | **0.66** | 0.53 | 0.04 | 0.37 |
| S3 | $h_3$ | 0.04 | **0.58** | -0.54 | -0.50 | -0.17 | 0.31 |
| S4 | $h_3$ | -0.05 | -0.39 | **0.87** | 0.07 | 0.16 | 0.26 |
| S5 | $h_3$ | 0.32 | 0.27 | **0.80** | 0.27 | 0.11 | -0.31 |
| S6 | $h_3$ | 0.34 | **0.76** | 0.24 | -0.08 | 0.16 | 0.47 |
| L1 | $h_3$ | 0.45 | 0.03 | **0.83** | -0.33 | -0.02 | 0.09 |
| L2 | $h_3$ | 0.33 | 0.62 | 0.05 | **0.70** | -0.12 | -0.15 |
| L3 | $h_3$ | -0.28 | -0.32 | -0.24 | -0.11 | 0.45 | -0.75 |
| L4 | $h_3$ | -0.02 | -0.22 | 0.16 | -0.47 | -0.38 | **0.75** |
| L5 | $h_3$ | 0.06 | **0.63** | 0.53 | -0.20 | 0.51 | -0.14 |
| L6 | $h_3$ | 0.10 | 0.31 | -0.17 | -0.01 | 0.18 | **0.92** |

## Notes

https://doi.org/10.6084/m9.figshare.14913051.v3

### Competing Interest Statement

The authors have declared no competing interest.

### Summary of Updates

All analyses and results remain the same. The introduction and discussion have been re-written to be more explicit about the broader implications.

### Funding

- Australian Research Council

